# Lustro: High-throughput optogenetic experiments enabled by automation and a yeast optogenetic toolkit

**DOI:** 10.1101/2023.04.07.536078

**Authors:** Zachary P Harmer, Megan N McClean

## Abstract

Optogenetic systems use genetically-encoded light-sensitive proteins to control cellular processes. This provides the potential to orthogonally control cells with light, however these systems require many design-build-test cycles to achieve a functional design and multiple illumination variables need to be laboriously tuned for optimal stimulation. We combine laboratory automation and a modular cloning scheme to enable high-throughput construction and characterization of optogenetic split transcription factors in *Saccharomyces cerevisiae*. We expand the yeast optogenetic toolkit to include variants of the cryptochromes and Enhanced Magnets, incorporate these light-sensitive dimerizers into split transcription factors, and automate illumination and measurement of cultures in a 96-well microplate format for high-throughput characterization. We use this approach to rationally design and test an optimized Enhanced Magnet transcription factor with improved light-sensitive gene expression. This approach is generalizable to high-throughput characterization of optogenetic systems across a range of biological systems and applications.

## Introduction

Optogenetics is a powerful technique that allows for dynamic, spatial, and temporal control over cellular behavior using light^1–3^. Optogenetics leverages light-sensitive proteins, taking advantage of light responsive changes in protein conformation to actuate processes inside the cell^4,5^. Such tools have been used to activate specific signaling pathways^6–8^, repress and activate transcription^9–11^, control protein localization^12–15^, and induce protein degradation^16,17^. A common approach is to control a process of interest by fusing effectors to light-activated hetero- or homodimerizers to generate activity through proximity. For example, light-sensitive split transcription factors (TFs) are frequently generated by fusing one protein of an optical heterodimerizer pair to a DNA-binding domain (DBD) and the other to an activation domain (AD). This allows expression of the gene of interest (GOI) to be activated by inducing dimerization (and reconstitution) of the split TF using light^18,19^.

One of the challenges of using optogenetics is that both prototyping a construct for a given application, as well as identifying appropriate illumination conditions, represent significant bottlenecks^5,6^. To generate a functional optogenetic construct, many factors need to be tuned and tested including expression levels, linker lengths, and choice of components. A cloning toolkit can be used to rapidly develop prototype constructs, and a yeast optogenetic toolkit was recently developed^9,10^, but it contains a relatively small fraction of the existing repertoire of light-sensitive proteins. In addition, once constructs are created, a high-throughput method is needed to characterize their function and activity in response to light. Bioreactor-based techniques have been developed that allow real-time measurement of light-sensitive cultures^20–24^ but they have limited throughput. Several tools allow for individual programming of LEDs in a microwell plate format such as the LPA^25^, optoPlate^26^, and LITOS^27^ and enable higher throughput light-stimulation. However, these approaches still lack a method for high-throughput and rapid measurement of the optogenetic system response. The recent optoPlateReader^28^ partially solves this problem, but requires the use of many biological replicates to obtain reliable data and lacks access to liquid handling capabilities, important for performing certain assays or long-term experiments.

To accelerate optogenetic prototyping, in this work we use laboratory automation to enable high-throughput screening and characterization of optogenetic systems with frequent and reliable measurements. This enables more rapid optimization and subsequent application of optogenetic systems. We dub this technique Lustro, after the latin verb signifying movement, surveying, and illumination. We couple this laboratory automation technique for high-throughput optogenetics stimulation and readout with a modular cloning toolkit^9,29,30^ to build and characterize a library of split transcription factors in the important biotechnology model organism budding yeast, or *Saccharomyces cerevisiae*.

## Results

### Integration of laboratory automation and light stimulation for high-throughput optogenetics

In order to increase testing throughput and reliability we developed an automated platform, Lustro, for screening and characterizing optogenetic systems. Lustro comprises an illumination device, a shaking device, and a plate reader integrated into a Tecan Fluent Automated Workstation (Figure 1a). A Robotic Gripper Arm (RGA) is able to move a microwell plate between these devices according to a programmed schedule. In our experiments, *S. cerevisiae* cultures are diluted into a 96-well plate with conditions measured in triplicate (Supplemental Figure 1). This plate is placed on the illumination device (an optoPlate) for 26.5 minutes to receive light induction by individually programmable LEDs. The Robotic Gripper Arm then moves the plate to the shaking device to shake at 1000 rpm for 60 seconds in order to resuspend the yeast cells. This ensures accurate and consistent measurements of a homogenous culture and also improves growth conditions. The plate is then moved to the plate reader to measure optical density and fluorescence before being moved back to the illumination device. The cycle is repeated on 30-minute intervals. Due to its size and weight, the optoPlate could not be incorporated onto the small plate shaker. We therefore shook cells intermittently, which led to a small but measurable lag in growth (Supplemental Figure 2).

**Figure 1.**
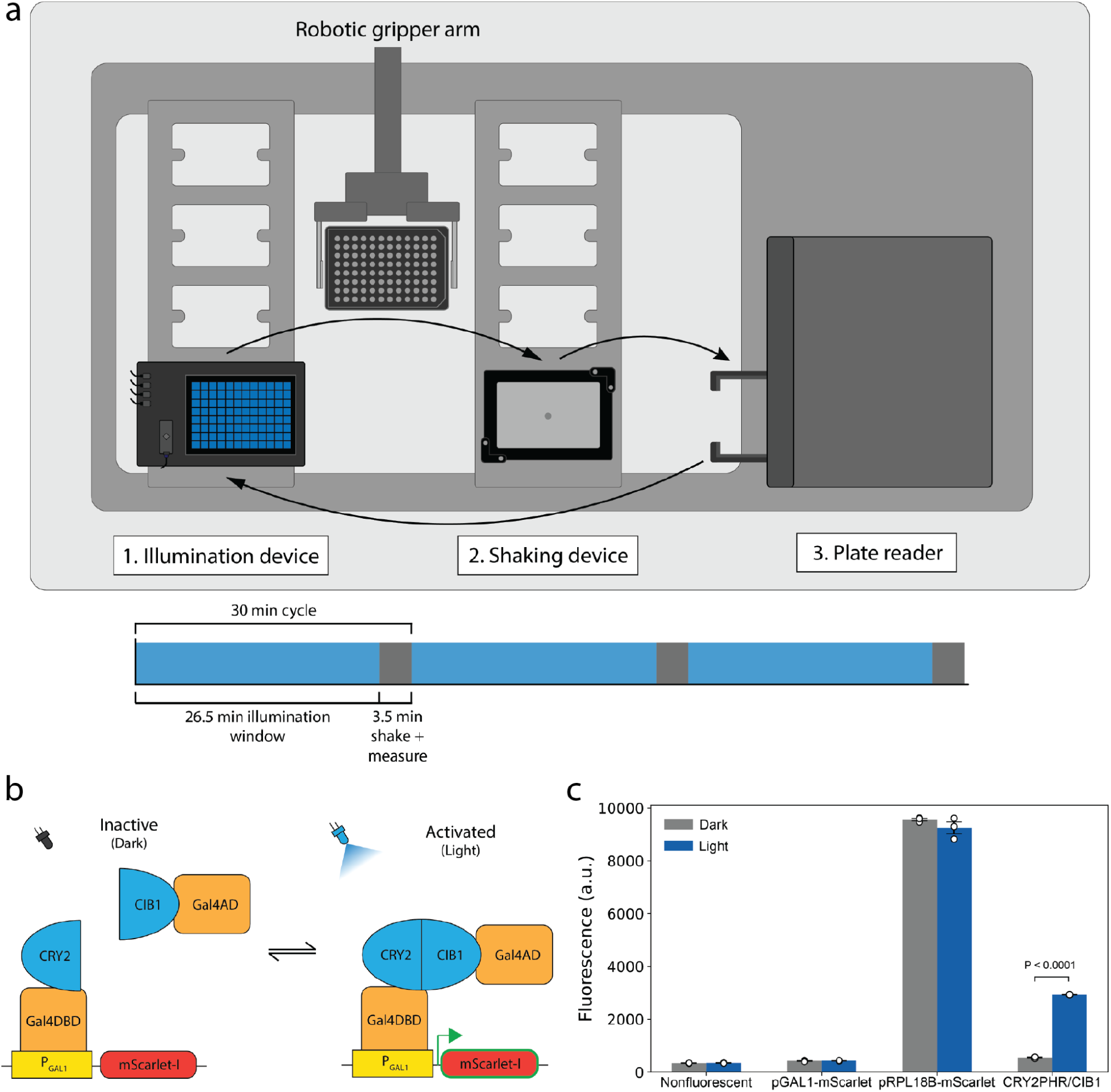
(a) The automated platform, Lustro. The Robotic Gripper Arm transfers microwell plates between the illumination device, shaking device, and plate reader. (b) Optogenetic split TF with CRY2 and CIB1 as the optical dimerizer pair. When Gal4AD is recruited to Gal4DBD (bound to pGAL1), expression of the gene of interest (mScarlet) is induced. (c) mScarlet fluorescence driven by induction of the CRY2PHR/CIB1 TF (p=0.000039, t-statistic=160, paired Student’s t-test), with non-fluorescent (negative) cells, a non-inducing control with only the reporter construct, and mScarlet under constitutive (pRPL18B) expression. Measurements were taken every 30 minutes and raw fluorescence values are shown for the light and dark conditions 10 hours into induction.

We demonstrated that the Lustro platform can be used to measure the activity of optogenetic split TFs (Figure 1b). These split TFs utilize the Gal4 DNA-binding domain (GAL4DBD) and Gal4 activation domain (Gal4AD) to drive expression of the fluorescent reporter mScarlet-I^31^ (hereafter referred to as mScarlet) from the pGAL1 promoter under light induction. This signal is readily measured by the plate reader. We induced a strain carrying a split transcription factor based on an optogenetic heterodimerizer pair (the cryptochrome CRY2PHR and its binding partner CIB1^19^) (Figure 1c) and compared it to a non-fluorescent strain (negative), a strain with only the reporter construct (pGAL1-mScarlet), and a strain with constitutive expression of the fluorescent reporter (pRPL18B-mScarlet).

### Combining optoPlate programming with automation allows for high-resolution time-course experiments

A powerful advantage of Lustro is the ability to easily record output over time (Figure 2a). This can reveal dynamic changes which would not be observed using a single end point measurement. In addition, it is known that culture growth stage and saturation can affect optogenetic systems differently^32^. We measured gene expression (using the fluorescent reporter) induced by a split TF (consisting of a cryptochrome variant, CRY2(535)/CIB1)^33^ every 30 minutes for 16 hours (Figure 2a). Measurements were taken every 30 minutes and reveal behavior of the optogenetic system in response to different light induction programs as the cell culture reaches saturation (Supplemental Figure 3).

**Figure 2.**
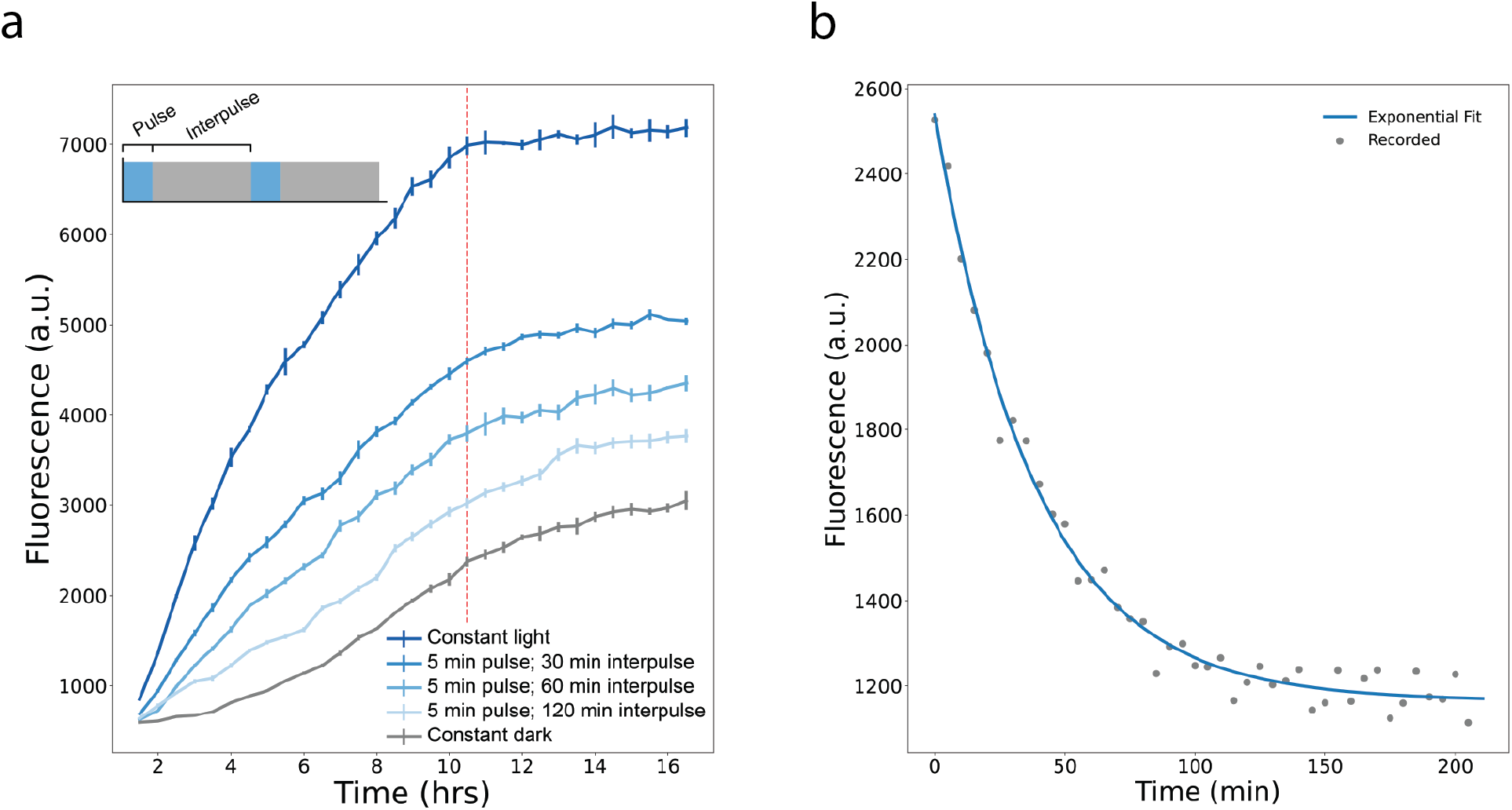
(a) Induction of a CRY2(535)/CIB1 split TF strain over time by light regimes with constant on times and varied off times. Each intermediate induction regime shown has a 5-minute pulse of light followed by an interval of darkness as indicated on the figure legend for the duration of light induction (shown on the horizontal axis). The red vertical line indicates when the cultures reach saturation (Supplemental Figure 3). (b) Decay of mRuby2 photoactivation effect. The dots (gray) are averages of triplicate measurements (arranged by the time since removal of light induction, shown on the horizontal axis) collated from the plots shown in Supplemental Figure 4a. The line (blue) shows the exponential decay curve fit to the data.

In preliminary experiments, we created strains using mRuby2^34,35^ as a fluorescent reporter (later replaced with mScarlet). However, a strain with mRuby2 under constitutive expression (pRPL18B) was unexpectedly found to temporarily exhibit higher fluorescence following light induction (Supplemental Figure 4). We were able to observe the kinetics of this photoactivated effect by using the automated platform, which we would not have been able to observe by taking single measurements at a delayed endpoint. The short sampling time (3.5 minutes to shake and measure a plate) and the ability to program illumination means that measurements can be taken with even finer timescale resolution if duplicate wells are used and illumination patterns are staggered. We used this approach to measure the timescale of the decay rate of the mRuby2 photoactivated effect. A strain constitutively expressing mRuby2 (pRPL18B-mRuby2) was induced with blue light and the timing of the light switching off between duplicate wells was staggered by 5-minute intervals so that measurements recorded on 30-minute intervals could show finer granularity (Supplemental Figure 4b). These measurements were combined and fit to an exponential decay curve (Figure 2b) and the half-life of this photosensitive effect was found to be 26.5 minutes. This effect may be related to the pH sensitivity of mRuby^35,36^. While frustrating for measuring the response of blue-light stimulated optogenetic systems, this effect could potentially be leveraged for other applications. For instance, to track protein movement and localization by stimulating mRuby2 in a defined location and observing the change and movement of the photosensitive fluorescence effect.

### A yeast optogenetic toolkit (yOTK) combined with Lustro allows for rapid prototyping and testing of optogenetic systems

Optogenetic split transcription factors can, in theory, be built from any pair of optically dimerizing proteins. However, these proteins have different properties, including their light sensitivity, photocycle kinetics, as well as their sensitivity to protein fusion and context^5^. In order to compare how different optical dimerizers tune the activity of optogenetic split TFs, we introduced new light-sensitive dimerizers as parts into the yeast optogenetic toolkit (yOTK)^9,29,30^. We specifically introduced several different cryptochrome^37^ variants (CRY2FL/CIB1^19^, CRY2PHR/CIB1^19^, CRY2(535)/CIB1^33^) and Enhanced Magnets (eMags) (eMagA/eMagB, eMagAF/eMagBF)^38^, selected to have different photocycle kinetics between light and dark states and light sensitivity (Table 1; see Supplemental Table 3 for full list of plasmids generated in this work). Using the toolkit, these optical dimerizer pairs are cloned into the same cellular context (Figure 3a) using Golden Gate assembly^39^ to rapidly and reliably assemble individual “part” plasmids into “cassette” plasmids (Supplemental Figure 5) containing split TFs that use the Gal4 activation domain (Gal4AD), the Gal4 DNA-binding domain (Gal4DBD), with the Gal1 promoter (pGAL1) driving expression of the fluorescent protein mScarlet. Cassettes containing the individual TF components (DBD, AD, or pGAL1) are further assembled into “multigene” plasmids for transformation into yeast.

**Figure 3.**
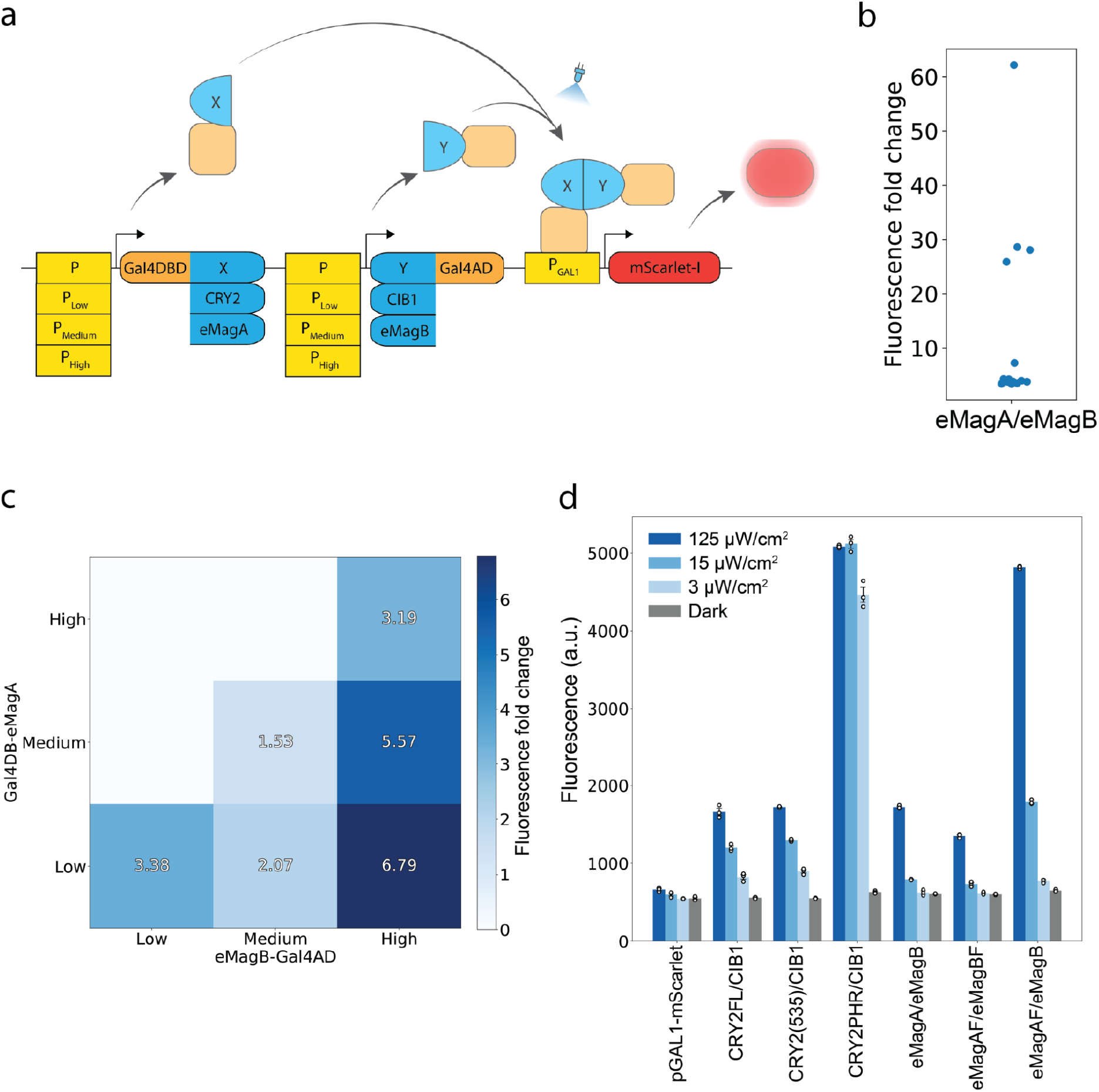
(a) Using the yeast optogenetic toolkit, multigene cassettes containing optical dimerizers fused to appropriate effector domains and controlled by a range of promoter strengths are created and integrated into the yeast genome.. Once expressed, the light-inducible split TF induces expression of the fluorescent reporter, pGAL1-mScarlet, in a light-dependent manner. (b) Each dot represents the fold change in mScarlet fluorescence between light and dark conditions for a different transformant of the eMagA/eMagB split TF. Data shown represent averaged triplicates measured after 12 hours of light induction. (c) Heat map showing fold change in fluorescence after 12 hours of induction between light and dark conditions for eMagA/eMagB split TF strains with components expressed at different levels. Horizontal and vertical axes identify the strains with each split TF component under low (pRPL18B), medium (pHHF1), or high (pTEF1) expression. (d) Fluorescence values after 12 hours of light induction at the indicated light intensities for strains expressing split TFs using the indicated protein pairs and a reporter-only pGal1-mScarlet control strain.

**Table 1.** Optogenetic parts added to the yeast MoClo toolkit in this work. Additional plasmid information found in Supplemental Table 3.

| **Dimerizer variant** | **Binding partner** | **Description** |
| --- | --- | --- |
| eMagA | eMagB, eMagBF, or eMagBM | Enhanced magnet dimerizer<sup>38</sup> |
| eMagAF | eMagB, eMagBF, or eMagBM | Enhanced magnet dimerizer with faster kinetics<sup>38</sup> |
| eMagB | eMagA or eMagAF | Enhanced magnet dimerizer<sup>38</sup> |
| eMagBF | eMagA or eMagAF | Enhanced magnet dimerizer with faster kinetics<sup>38</sup> |
| eMagBM | eMagA or eMagAF | Enhanced magnet dimerizer with slower kinetics (Figure 4) |
| CRY2FL | CIB1 | Full length CRY2<sup>19</sup> |
| CRY2PHR | CIB1 | CRY2 truncation (residues 1-498 of CRY2)<sup>19</sup> |
| CRY2(535) | CIB1 | CRY2 truncation (residues 1-535 of CRY2)<sup>33</sup> |

We used Lustro to test various induction programs and screen several colonies from each construct transformation. Different transformants of the same construct were found to have variable fold-change gene expression response to induction (Figure 3b). We hypothesize these differences are due to copy number integration variation, as has been seen previously^40^. Lustro allows for 46 transformants to be screened in each 96-well plate (light and dark conditions of each transformant alongside blank and negative controls, see Supplemental Figure 1), providing a robust and reliable method for identifying transformants with desired traits. For purposes of comparing the effects of different promoters and optical dimerizers, the lowest fold-change transformants were assumed to be single-copy integrations and selected. The transformants exhibiting a higher light-induced gene expression level, presumably due to multiple integrations, might be preferred for some applications and merit further exploration in a future study.

Tuning relative expression levels of the two components in split TF optogenetic systems is important for optimizing their activity^9^. We used the yeast optogenetic toolkit to develop strains with the DBD and AD components under different strength promoters and rapidly tested the strains with the Lustro automated platform. We generated light-inducible split TF strains with optical dimerizer eMagA and eMagB components^38^ under constitutive expression of low, medium, and high strength promoters (Figure 3c). Using Lustro, all construct transformants were screened and tested in two days. The strains use pRPL18B as the low promoter, pHHF1 as the medium, and pTEF1 as the high, based on characterizations done by Lee, et al 2015^29^.

We reasoned that excess expression levels of the DBD component relative to the AD component expression levels would result in suboptimal activation as there are limited binding sites in the genome and unbound DBD could sequester the AD component away from the DNA without providing gene expression activity. This effect has been seen in previous studies^9^, which demonstrated that higher expression of the DBD component relative to the AD component suppressed light-induced gene expression. Thus we only generated strains where the expression of the AD component is equal to or greater than the expression of the DBD component. For each expression strength of the AD component, a higher fold change in fluorescence corresponded to a lower expression strength of the DBD component, with the largest effect occurring for the AD component under highest expression and the DBD component under lowest expression (Figure 3c, lower right).

Different optical dimerizers are known to exhibit different light sensitivity^5,19,33,38,41,42^. The yOTK was used to generate strains with different optical dimerizers cloned into a similar split TF context and Lustro was used to characterize the light sensitivity of these strains (Figure 3d). We found that TFs using CRY2FL and CRY2(535) exhibited similar levels of sensitivity to light intensity, while the CRY2PHR TF exhibited very high sensitivity to even low doses of constant light. CRY2PHR is a truncation of CRY2FL that exhibits both higher basal and light-induced activity^19^. CRY2(535) is an intermediate-length truncation that produces intermediate activation and background, as compared to CRY2FL and CRY2PHR^33^. Comparatively, TFs using the Enhanced Magnets (detailed below) exhibited less sensitivity to low levels of light intensity. TFs using a variant of eMagA/eMagB designed to have faster photocycle kinetics, eMagAF/eMagBF^38^, had a lower gene expression response than TFs using eMagA/eMagB, but similar light sensitivity. Surprisingly, a split TF using a combination of the two, eMagAF/eMagB, exhibited a much higher gene expression level (and somewhat higher light sensitivity) than TFs using either eMagA/eMagB or eMagAF/eMagBF.

### Combining the yOTK and Lustro to generate an optimized Magnet-based split TF

The original Magnet proteins were developed by introducing positively and negatively charged residues into the Ncap homodimer interface of the homodimerizer protein Vivid, a naturally occurring light-sensitive protein in *Neurospora crassa*^43^. Subsequent mutations were introduced to reduce (pMagFast2) or increase (nMagHigh1) reversion time to the dark state. The Enhanced Magnets eMagA and eMagB used in this study were generated from nMagHigh1 and pMagFast2 (respectively) by introducing mutations to improve thermal stability and binding activity^38^. To develop a version of eMagB that reverts to the dark state more slowly, we introduced mutations into another Magnet protein, pMagFast1 (a slower reverting version of pMagFast2), generating an “Enhanced” pMagFast1 protein, eMagBM (Figure 4a).

**Figure 4.**
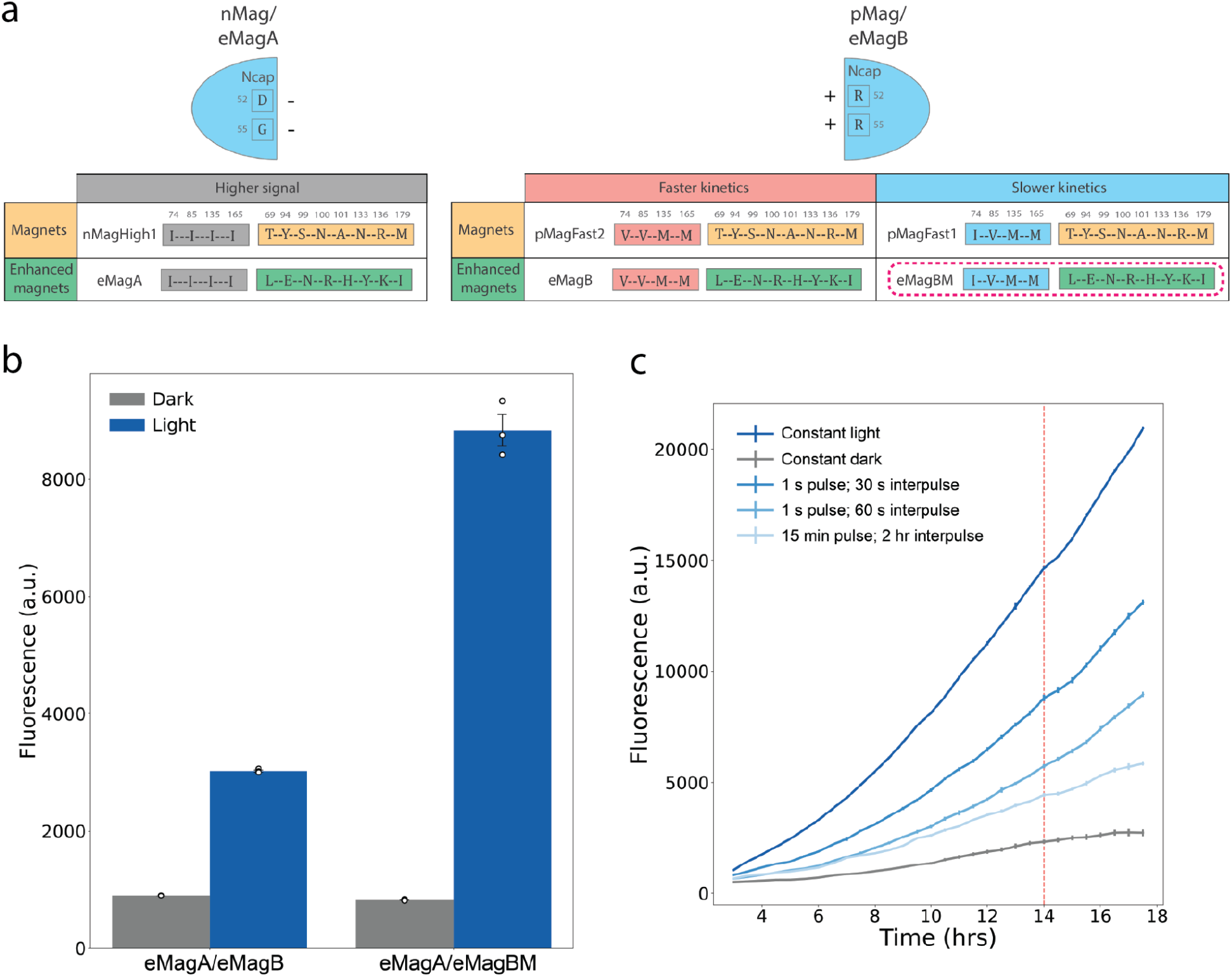
(a) Table showing the mutations made to design eMagBM (highlighted in pink dashed line). Mutation of residue 74 to I was associated with slower kinetics in pMag. (b) Fluorescence values are shown after 12 hours of light induction between light and dark samples of strains with eMagB- or eMagBM-based split TFs. Fluorescence with the eMagA/eMagB split TF system increases 3.4 fold under light induction and fluorescence of the eMagA/eMagBM split TF system increases 10.8 fold. All split TF components are expressed under pRPL18B. (c) Fluorescence is shown for the eMagA/eMagBM TF strain under different induction regimes that vary the pulse on and pulse off times in minutes and seconds. The vertical red line shows when the cultures reach saturation, and the horizontal axis shows time since the start of induction.

We used Lustro to characterize split TFs with eMagBM and compare them to split TFs using eMagB. Induction with eMagBM-based TFs was found to be higher than induction with eMagB-based TFs as we had anticipated (Figure 4b) and was tunable by varying light pulse and interpulse duration (Figure 4C). Interestingly, induction with the Magnet-based TFs was found to continue to increase fluorescence even after cultures reached saturation at around 12 hours (data shown for eMagA/eMagBM in Figure 4c), which might be useful for high cell density bioproduction schemes^44^. This presents a contrast to the activity of the CRY2/CIB1-based TFs (data shown for CRY2PHR/CIB1 in Figure 2a), where fluorescence plateaus around the saturation point at 10 hours.

To further optimize the eMagA/eMagBM split TF, we cloned constructs with the components of the eMagBM-based split TF system each expressed under different promoter strengths, as was done with the eMagA/eMagB split TF in Figure 3c. Transformants were screened as in Figure 3b to identify single-copy integrations for comparison (Supplemental Figure 6). Different expression levels of the eMagA/eMagBM split TF exhibited a similar pattern to the expression levels of the eMagA/eMagB split TF shown in Figure 3c, but with a higher expression of the reporter at all component expression levels (Supplemental Figure 7).

## Conclusion

Optogenetics is a potent tool for the precise control of biological activity. Combining advancements in high-throughput strain construction with rapid screening creates a pipeline to improve the speed and robustness of design-build-test cycles. In this study, we developed an automated high-throughput platform for optogenetics in microwell plates, Lustro, that enables rapid screening and comparison of different optogenetic systems. With the Lustro automated platform, it is possible to quickly test and optimize different light-sensitive proteins for their desired purposes. We demonstrate that Lustro can be used for growing and inducing cells with only a small lag in growth due to shaking and temperature conditions. Lustro is able to quickly screen transformants, allowing for selection of strains with desired properties. The combination of programmed light to stagger light conditions between duplicate wells allows for rapid phenomena to be measured, as demonstrated by measuring the decay rate of mRuby2 photoactivation. We combine Lustro with a modular cloning toolkit^9,29,30^ to create a pipeline that allows for testing and tuning the design of different optogenetic systems and comparing the response of different optogenetic systems to various light induction conditions. We use Lustro to characterize a split TF that uses a new, rationally designed Enhanced Magnet^38,41^ with a higher level of gene expression and expanded that cloning toolkit to include more optogenetic tools.

The Lustro platform is highly adaptable and can be generalized to work with other laboratory automation robots, illumination devices, plate readers, cell types, and optogenetic systems, including those responsive to other wavelengths of light^18,45^. For example, the optoPlate could be exchanged with the LPA^25^, LITOS^27^, or optoWELL-24^46^. It can also be adapted to perform any assay that can be done using a plate reader or a pipetting robot, which expands its potential applications. Laboratory automation has long been a staple in the pharmaceutical industry and genomics, as it dramatically increases throughput and frees up researchers from repetitive tasks to perform higher-level analysis^47,48^. Recent years have seen an increase of automation in optogenetics experiments^22,49^, and performing automated experiments in a microwell plate format increases throughput and allows for integration with other types of assays. The strategy of rapidly prototyping optogenetic circuit construction strategies in a microwell plate format complements other strategies for scaling up production with optogenetics^23^. Lustro can be further modified to facilitate automated dilutions for continuous culture applications, which is advantageous for long-term experiments. The possibility of frequent measurements allows Lustro to be adapted for cybergenetic feedback control, using real-time feedback and adjustments to alter the experimental conditions^6,49,50^. While all of the experiments performed here used the Gal4DBD for consistency, these split TFs can be designed using other DBDs or a targetable deactivated nuclease (such as dCas9^10^) to allow for screening of dynamic expression changes in multiple genes, making it a highly valuable tool in functional genomics studies^10,51^. Our automated high throughput platform Lustro offers a highly versatile and adaptable tool for rapidly screening and optimizing optogenetic systems, which will enable many new avenues of exploration into dynamic gene expression control.

## Materials and Methods

### Strains, Media, and Culture Conditions

Single-construct strains were assembled by transforming NotI-digested multigene plasmids into BY4741^52^ *Saccharomyces cerevisiae* MATα HIS3D1 LEU2D0 LYS2D0 URA3D0 GAL80::KANMX GAL4::spHIS5. Transformations were performed according to an established LiAc/SS carrier DNA/PEG protocol^53^. Constructs were genomically integrated to reduce cell-to-cell variability. Integrations were done at the URA3 site and transformants were selected using SC-Ura dropout media.

Yeast strains were inoculated from colonies on a YPD agar^54^ plate into 3mL liquid SC media^55^ and grown overnight at 30°C, shaking. Overnight cultures were diluted to OD_700_ = 0.1 in SC media. 200 μL of each culture was then added to each well of a 96-well black-wall, glass-bottom plate (Cat. #P96-1.5H-N). OD_700_ was used to avoid bias from the red fluorescent mScarlet-I^31,56^. All strains and conditions were measured in triplicate after initial transformant screening.

Cloning was carried out using a modular cloning toolkit as previously described^9,29,30^. In brief, part plasmids were constructed using BsmBI Golden Gate assembly of PCR products (primers in supplemental) or gBlocks (sequences in supplemental) into the part plasmid entry vector (yTK001). Part plasmids were subsequently assembled into cassette plasmids using BsaI Golden Gate assembly. Cassette plasmids were assembled into multigene plasmids using BsmBI Golden Gate assembly.

### optoPlate Configuration and Calibration

The optoPlate for light induction was constructed and calibrated according to previously published methods^26,57^. Re-calibration of the optoPlate was found to be necessary for consistent illumination since the time of its initial calibration by Grødem et al^57^ (possibly due to decay of the LEDs). The optoPlate was programmed for each experiment using scripts found here: https://github.com/mccleanlab/Optoplate-96. An intensity of 125 μW/cm^2^ was used for all experiments unless otherwise specified.

### Automated characterization of optogenetic systems on the Tecan Fluent Automation Workstation and Spark plate reader

Automated experiments were carried out on a Tecan Fluent Automation Workstation, programmed using the Tecan Fluent Control visual interface software. The Fluent is equipped with an optoPlate^26^, BioShake 3000-T elm heater shaker for well plates, a Tecan Spark plate reader, and a Robotic Gripper Arm (RGA) for moving plates and plate lids. Experiments were done using Cellvis 96-well glass bottom plates with #1.5 cover glass (Cat. # P96-1.5H-N). Plates are covered with a lid for all experiments except for those measuring the photoactivation effect of mRuby2^34,35^. For experiments measuring the photoactivation effect of mRuby2, the plate was covered in a Breathe-Easy polyurethane sealing membrane (Diversified Biotech, BEM-1) because these conditions created a stronger light-induced fluorescent signal. Each 96-well plate with culture diluted to OD_700_ = 0.1 was incubated for 5 hours in the dark at 30°C, shaking, before beginning light induction. For each light induction experiment, the plate was placed on the optoPlate for 26.5 minutes at 21°C, transferred to the BioShake to shake at 1000 rpm for 1 minute, and then transferred to the Tecan Spark plate reader to read optical density and fluorescence (with the lid temporarily removed by the Robotic Gripper Arm to ensure accurate OD readings). The plate is then transferred back to the optoPlate, and this cycle is repeated over the course of the experiment. Optical density is measured at 700 nm to avoid bias from measuring red fluorescent mScarlet-I^31,56^. For mScarlet-I, fluorescence is measured with excitation at 563 nm and emission at 606 nm, with Z=28410, and an optical gain of 130.

### Data analysis

Error bars shown represent the standard error of sample means performed in technical triplicate. Fold change shown is the raw fluorescence value of the induced strain divided by the raw fluorescence of the dark control. Exponential decay of mRuby2 photoactivation measurements were fit to the decay curve y=a.e^-b.x^+c using the curve_fit function from the scipy.optimize package in Python^58^.

## Materials Availability

Key plasmids have been deposited on Addgene. For all other reagent requests, please contact the corresponding author.

## Acknowledgments

This work was supported by National Institutes of Health grant R35GM128873 and National Science Foundation grant 2045493 (awarded to M.N.M.). Flow cytometry was enabled by the University of Wisconsin Carbone Cancer Center Support Grant P30 CA014520. Megan Nicole McClean, PhD holds a Career Award at the Scientific Interface from the Burroughs Wellcome Fund. Z.P.H. was supported by an NHGRI training grant to the Genomic Sciences Training Program 5T32HG002760. We thank Amit Nimunkar and Edvard Grødem for building and modifying the optoPlate. We acknowledge fruitful discussions with McClean lab members and in particular we are grateful to Neydis Moreno and Kevin Lauterjung for providing comments on the manuscript and to Stephanie Geller for providing pMM0773.

## Author Contributions

Z.P.H. and M.N.M conceived of the study; Z.P.H. designed optogenetic parts, developed the Lustro platform, performed experiments, and analyzed data. M.N.M. provided funding. Z.P.H. wrote the original draft of the manuscript and both Z.P.H. and M.N.M wrote, reviewed, and edited the final manuscript.

