## Supplemental Material for "Lustro: High-throughput optogenetic experiments enabled by automation and a yeast optogenetic toolkit"

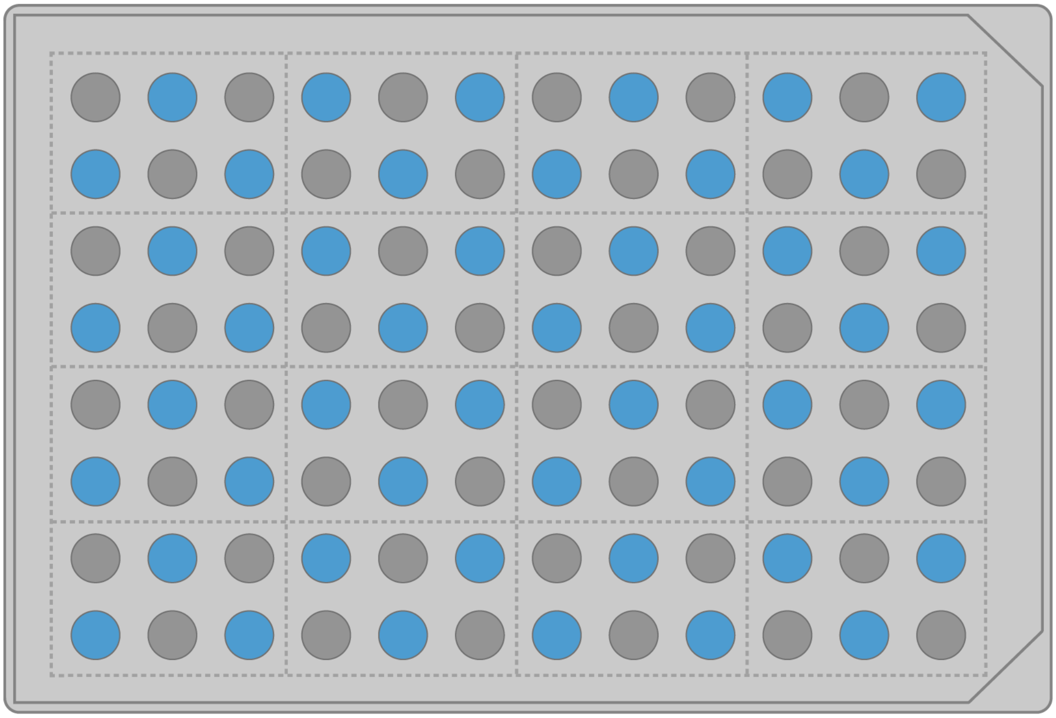

**Supplemental Figure 1**. Sample 96-well plate layout. Light conditions are checkerboarded (blue wells correspond to light induction, and gray wells correspond to dark treatment). Dotted lines indicate groupings of triplicate samples (with light and dark conditions) for each strain.

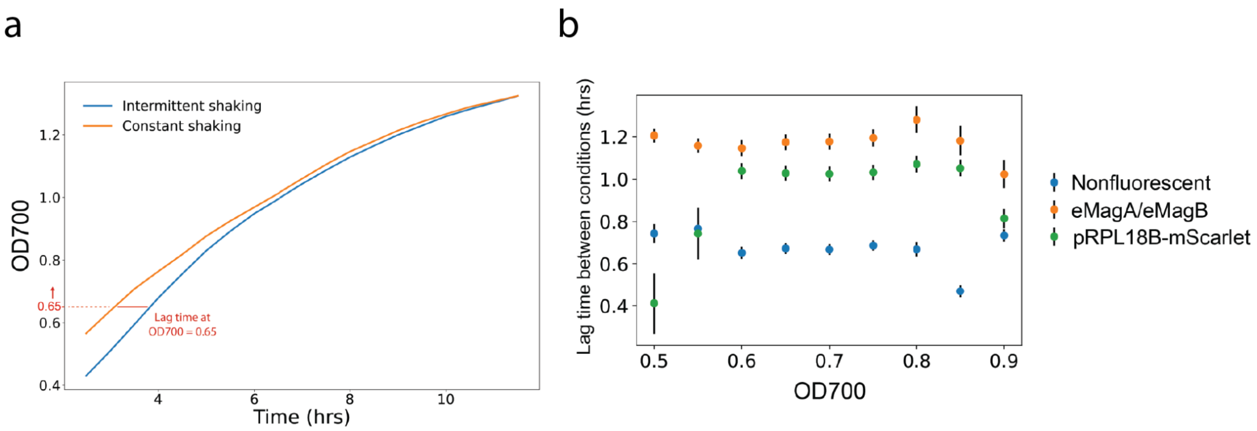

**Supplemental Figure 2.** (a) Diagram showing growth in continuously shaking conditions relative to intermittent shaking measured using OD700. The lag was measured at OD700=0.7 based on the sensitivity analysis in (b). Lag time is measured by subtracting the time needed for constantly shaking samples to reach certain OD700 values from the time needed for intermittently shaking samples to reach the same density. Measurements are taken every 30 minutes, and values are interpolated by linear approximation between measurements. (b) Sensitivity analysis of lag time to reach certain OD700 values between intermittent and constant shaking conditions for non-fluorescent (negative), eMagA/eMagB, and reporter-only (pRPL18B-mScarlet) strains. Lag is constant between OD700 of 0.6 and 0.8, therefore we quantified lag at OD700=0.7 for each strain (non-fluorescent: 0.67 hrs, eMagA/eMagB: 1.18 hrs, pRPL18B-mScarlet: 1.02 hrs).

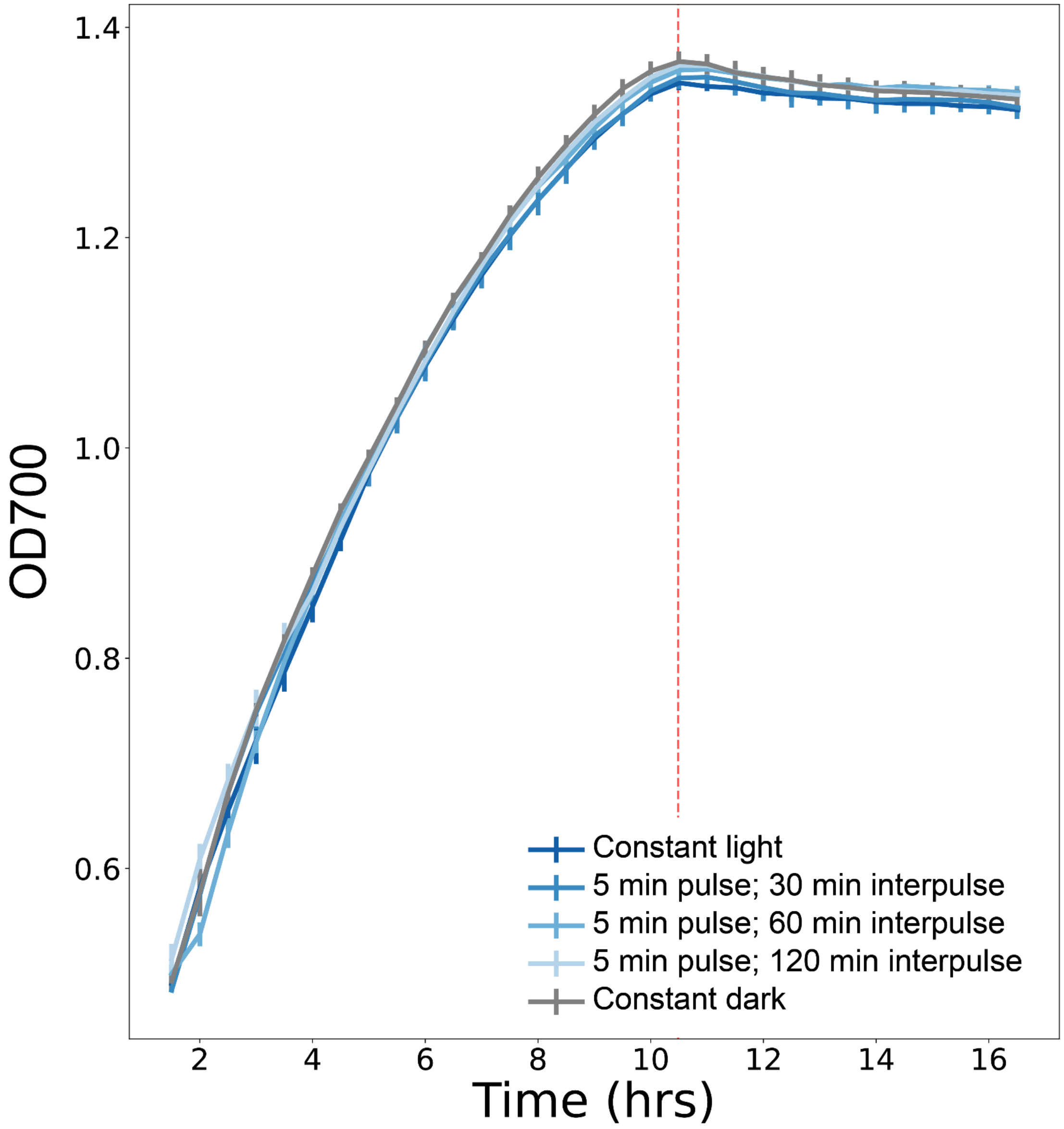

**Supplemental Figure 3.** OD700 values for the CRY2(535)/CIB1 split TF strain used in Figure 2a. The vertical red line indicates when the cultures reach saturation. There were no significant differences in OD700 between cultures exposed to different light induction programs.

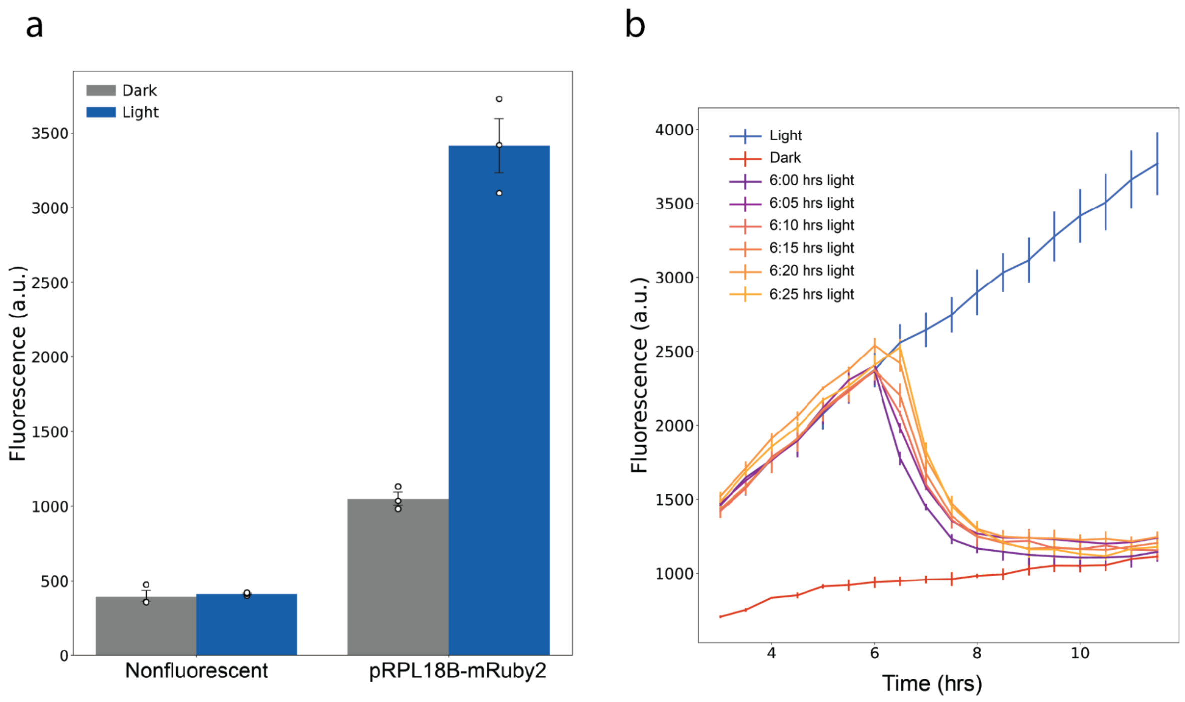

**Supplemental Figure 4**. (a) Photostimulation of the pRPL18B-mRuby2 strain. Lengths of light induction are shown in the legend; time since beginning light induction is shown on the horizontal axis. (b) Induction of a non-fluorescent control strain (negative) and the pRPL18B-mRuby2 strain after 12 hours. The photoactivated mRuby2 strain shows significantly higher fluorescence than the dark control (Student's t-test; t-statistic: 11.77, p-value: 0.007).

| Assembly Connector | Promoter | Coding Sequences | | Terminator | Assembly Connector | *S. cerevisiae* marker | 3' Homology | *E. coli* marker and origin | 5' Homology |
|---|---|---|---|---|---|---|---|---|---|
| 1 | 2 | 3 | | 4 | 5 | 6 | 7 | 8a | 8b |
| ConLS | pRPL18B | Gal4AD-CIB1 | | tENO1 | ConR1 | URA3 | URA3 3' Hom | KanR-ColE1 | URA3 5' Hom |
| ConL1 | pHHF1 | mScarlet-I | | | ConR2 | | | | |
| ConL2 | pTEF1 | 3a | 3b | | ConRE | | | | |
| | pGAL1 | Gal4DBD | Gal4AD | | | | | | |
| | | eMagB | CRY2FL | | | | | | |
| | | eMagBF | CRY2PHR | | | | | | |
| | | eMagBM | CRY2(535) | | | | | | |
| | | | eMagA | | | | | | |
| | | | eMagAF | | | | | | |

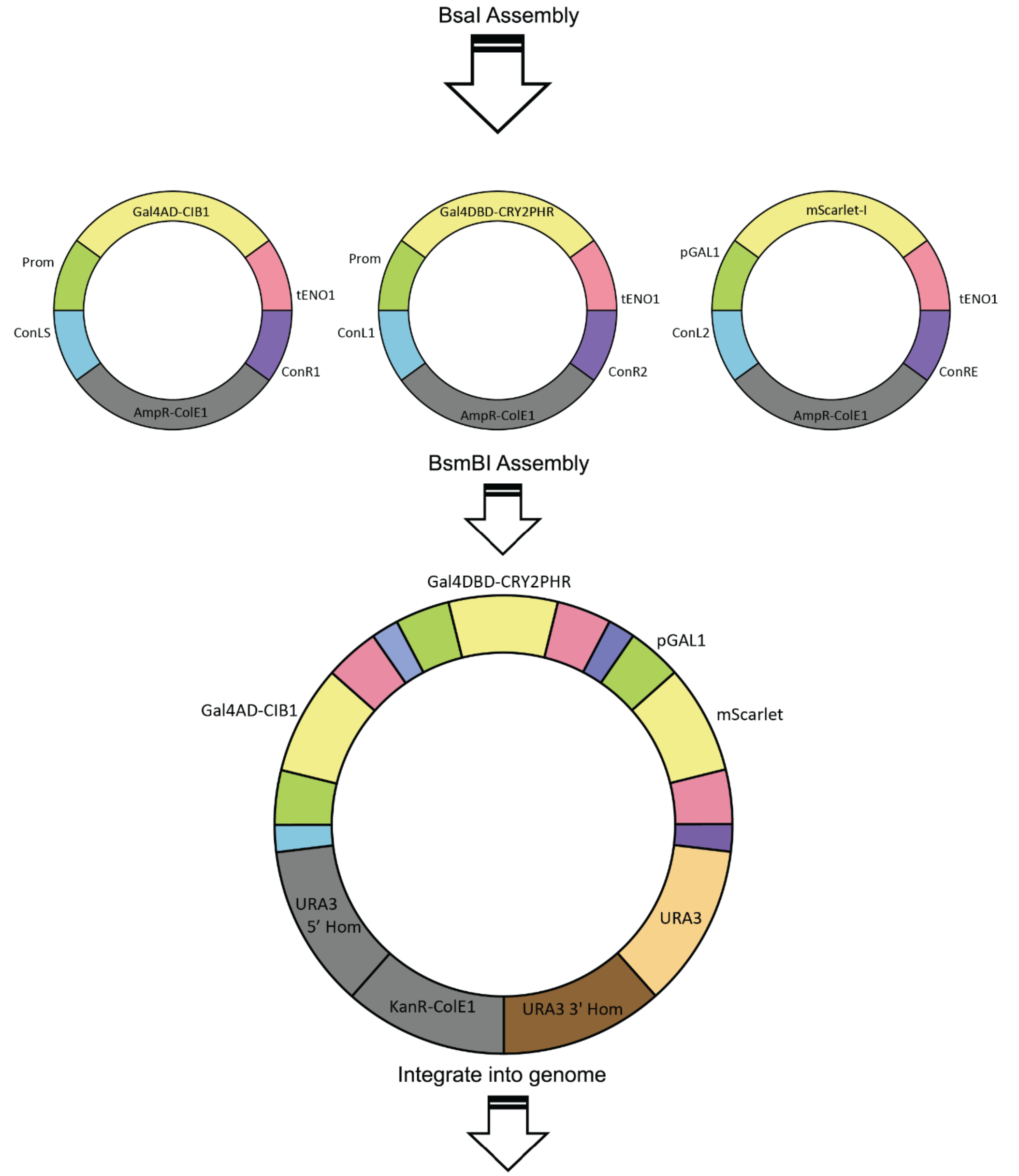

**Supplemental Figure 5**. Diagram showing how toolkit parts are assembled to form individual cassettes, which are further assembled into multigene cassettes for genome integration according to previously published methods[1,2]. Figure colors and scheme are adapted for consistency from Lee, et al 2015[2].

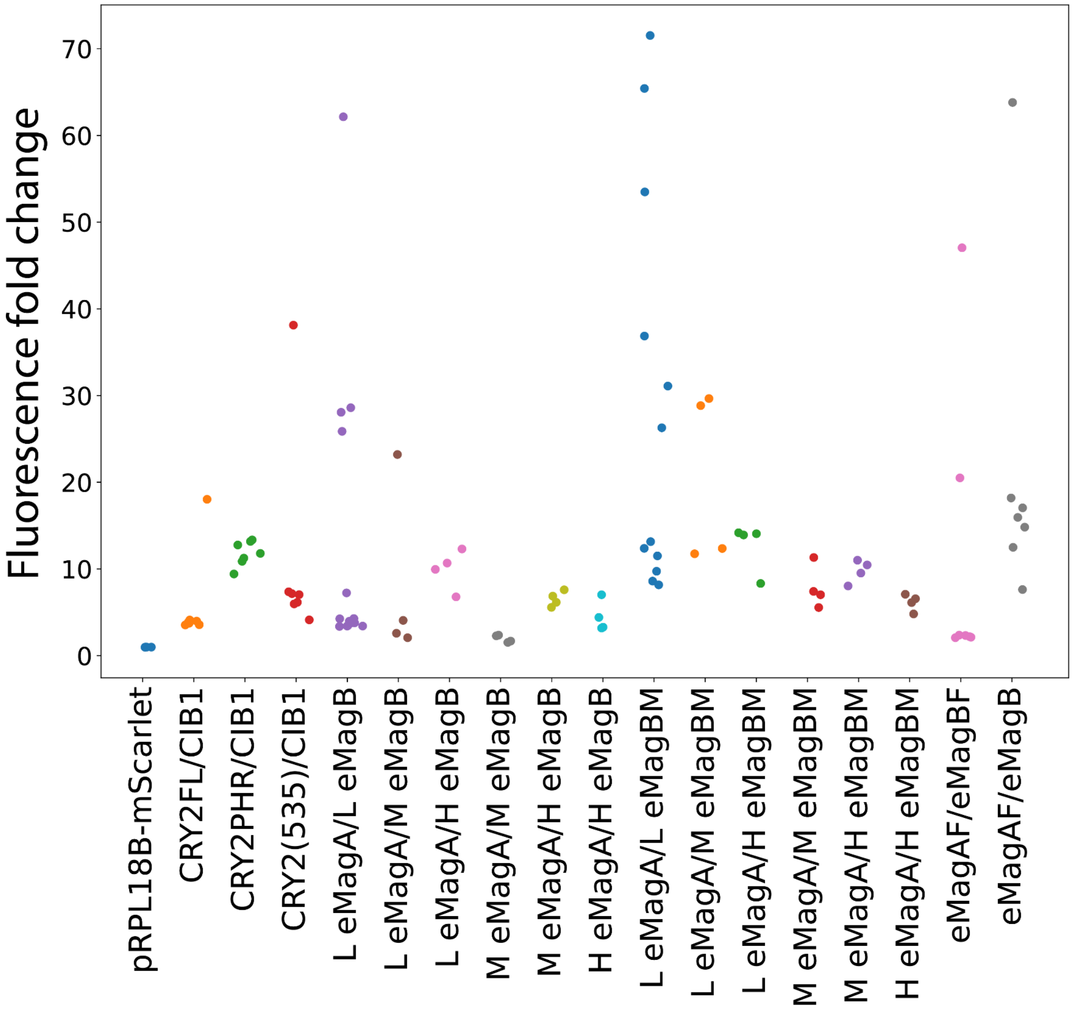

**Supplemental Figure 6**. Fluorescence fold-change between light and dark conditions for various construct transformants after 12 hours of light induction. L/M/H designations indicate whether each split TF component is under low (pRPL18B), medium (pHHF1), or high (pTEF1) expression[2].

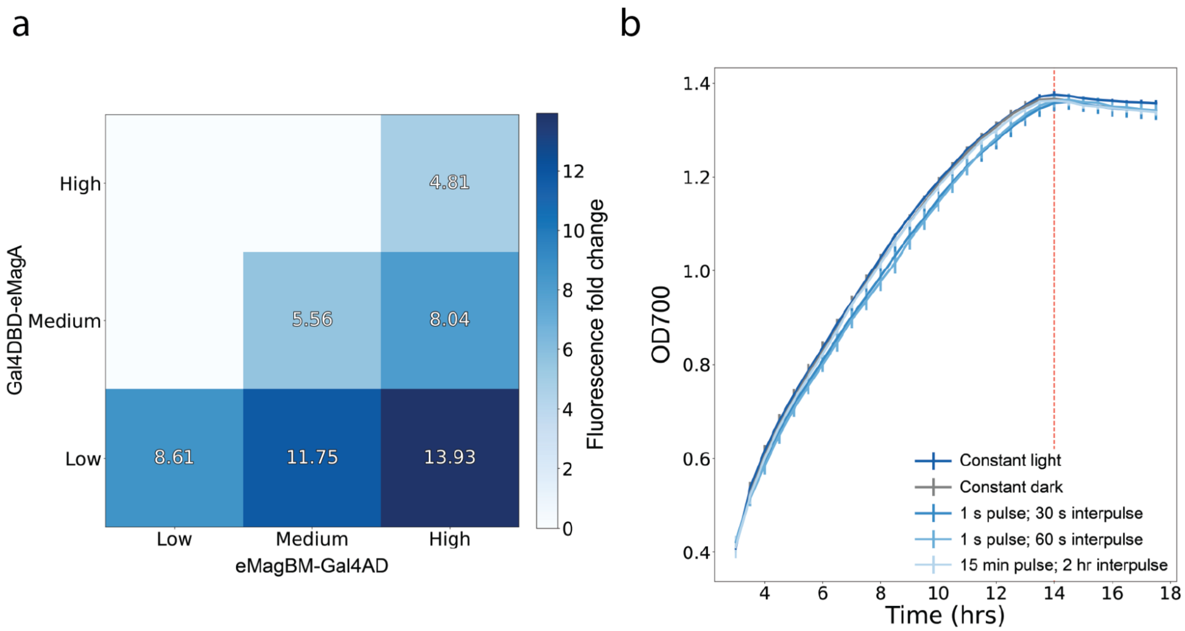

**Supplemental Figure 7.** (a) Heat map showing fold change in fluorescence after 12 hours of induction between light and dark conditions for eMagA/eMagBM split TF strains with components expressed at different levels. Horizontal and vertical axes identify the strains with each split TF component under low (pRPL18B), medium (pHHF1), or high (pTEF1) expression. Compare to eMagA/eMagB strains in Figure 3c. (b) OD700 values for the eMagA/eMagBM split TF strain used in Figure 4c. The vertical red line indicates when the cultures reach saturation. There were no significant differences in OD700 between cultures exposed to different light induction programs.

### Supplemental Table 1: Yeast strains

| ID | Alias | Genotype | Description | Source |
|---|---|---|---|---|
| yMM1444 | mRuby2 MEDIUM | Mat alpha trp1Δ63 leu2Δ1::LEU2-pRPL18B-mRUBY2-tADH1 ura3-52 HO::S40NLS-VP16AD - CIB1 loxP-KlURA3-loxP SV40NLS-Zif268DBD-CRY2PHR | Benchmark for mRuby2 expression (pRPL18B) | An-adirekkun et al. 2020 Biotechnol Bioeng |
| yMM1715 | Ura int ctrl | BY Matα ura3Δ0::URA3-sfGFP his3D1 leu2D0 lys2D0 gal80::KANMX gal4::spHIS5 | Ura3 integration control | This work |
| yMM1728 | spacer, spacer, pGAL1-mScarlet-I | BY Matα ura3Δ0::5' Ura3 homology, spacer, spacer, pGAL1-mScarlet-I-tENO1, Ura3, Ura 3' homology his3D1 leu2D0 lys2D0 gal80::KANMX gal4::spHIS5 | pGAL1-mScarlet-I control | This work |
| yMM1729 | spacer, spacer, pRPL18B-mScarlet-I | BY Matα ura3Δ0::5' Ura3 homology, spacer, spacer, pRPL18B-mScarlet-I-tENO1, Ura3, Ura 3' homology his3D1 leu2D0 lys2D0 gal80::KANMX gal4::spHIS5 | pRPL18B-mScarlet-I control | This work |
| yMM1731 | pRPL18B-Gal4DBD-CRY2PHR, pRPL18B-Gal4AD-CIB1, pGAL1-mScarlet-I | BY Matα ura3Δ0::5' Ura3 homology, pRPL18B-Gal4DBD-CRY2PHR-tENO1, pRPL18B-Gal4AD-CIB1-tENO1, pGAL1-mScarlet-I-tENO1, Ura3, Ura 3' homology his3D1 leu2D0 lys2D0 gal80::KANMX gal4::spHIS5 | pRPL18B-CRY2PHR, pRPL18B-CIB1 | This work |
| yMM1733 | pRPL18B-Gal4DBD-CRY2, pRPL18B-Gal4AD-CIB1, pGAL1-mScarlet-I | BY Matα ura3Δ0::5' Ura3 homology, pRPL18B-Gal4DBD-CRY2-tENO1, pRPL18B-Gal4AD-CIB1-tENO1, pGAL1-mScarlet-I-tENO1, Ura3, Ura 3' homology his3D1 leu2D0 lys2D0 gal80::KANMX gal4::spHIS5 | pRPL18B-CRY2, pRPL18B-CIB1 | This work |
| yMM1734 | pRPL18B-Gal4DBD-eMagA, pRPL18B-eMagB-Gal4AD, pGAL1-mScarlet-I | BY Matα ura3Δ0::5' Ura3 homology, pRPL18B-Gal4DBD-eMagA-tENO1, pRPL18B-eMagB-Gal4AD-tENO1, pGAL1-mScarlet-I-tENO1, Ura3, Ura 3' homology his3D1 leu2D0 lys2D0 gal80::KANMX gal4::spHIS5 | pRPL18B-eMagA, pRPL18B-eMagB | This work |

| | | | | |
|---|---|---|---|---|
| yMM1760 | pRPL18B-Gal4DBD-eMagA, pTEF1-eMagB-Gal4AD, pGAL1-mScarlet-I | BY Matα ura3Δ0::5' Ura3 homology, pRPL18B-Gal4DBD-eMagA-tENO1, pTEF1-eMagB-Gal4AD-tENO1, pGAL1-mScarlet-I-tENO1, Ura3, Ura 3' homology KanR-ColE1 his3D1 leu2D0 lys2D0 gal80::KANMX gal4::spHIS5 | pRPL18B-eMagA, pTEF1-eMagB | This work |
| yMM1761 | pTEF1-Gal4DBD-eMagA, pTEF1-eMagB-Gal4AD, pGAL1-mScarlet-I | BY Matα ura3Δ0::5' Ura3 homology, pTEF1-Gal4DBD-eMagA-tENO1, pTEF1-eMagB-Gal4AD-tENO1, pGAL1-mScarlet-I-tENO1, Ura3, Ura 3' homology his3D1 leu2D0 lys2D0 gal80::KANMX gal4::spHIS5 | pTEF1-eMagA, pTEF1-eMagB | This work |
| yMM1763 | pRPL18B-Gal4DBD-CRY2(535), pRPL18B-Gal4AD-CIB1, pGAL1-mScarlet-I | BY Matα ura3Δ0::5' Ura3 homology, pRPL18B-Gal4DBD-CRY2(535)-tENO1, pRPL18B-Gal4AD-CIB1-tENO1, pGAL1-mScarlet-I-tENO1, Ura3, Ura 3' homology his3D1 leu2D0 lys2D0 gal80::KANMX gal4::spHIS5 | pRPL18B-CRY2(535), pRPL18B-CIB1 | This work |
| yMM1764 | pRPL18B-Gal4DBD-eMagAF, pRPL18B-eMagBF-Gal4AD, pGAL1-mScarlet-I | BY Matα ura3Δ0::5' Ura3 homology, pRPL18B-Gal4DBD-eMagAF-tENO1, pRPL18B-eMagBF-Gal4AD-tENO1, pGAL1-mScarlet-I-tENO1, Ura3, Ura 3' homology his3D1 leu2D0 lys2D0 gal80::KANMX gal4::spHIS5 | pRPL18B-eMagAF, pRPL18B-eMagBF | This work |
| yMM1765 | pRPL18B-Gal4DBD-eMagA, pRPL18B-eMagBM-Gal4AD, pGAL1-mScarlet-I | BY Matα ura3Δ0::5' Ura3 homology, pRPL18B-Gal4DBD-eMagA-tENO1, pRPL18B-eMagBM-Gal4AD-tENO1, pGAL1-mScarlet-I-tENO1, Ura3, Ura 3' homology his3D1 leu2D0 lys2D0 gal80::KANMX gal4::spHIS5 | pRPL18B-eMagA, pRPL18B-eMagBM | This work |
| yMM1769 | pRPL18B-Gal4DBD-eMagA, pHHF1-eMagBM-Gal4AD, pGAL1-mScarlet-I | BY Matα ura3Δ0::5' Ura3 homology, pRPL18B-Gal4DBD-eMagA-tENO1, pHHF1-eMagBM-Gal4AD-tENO1, pGAL1-mScarlet-I-tENO1, Ura3, Ura 3' homology his3D1 leu2D0 lys2D0 gal80::KANMX gal4::spHIS5 | pRPL18B--eMagA, pHHF1-eMagBM | This work |
| yMM1770 | pRPL18B-Gal4DBD-eMagA, pTEF1-eMagBM-Gal4AD, pGAL1-mScarlet-I | BY Matα ura3Δ0::5' Ura3 homology, pRPL18B-Gal4DBD-eMagA-tENO1, pTEF1-eMagBM-Gal4AD-tENO1, pGAL1-mScarlet-I-tENO1, Ura3, Ura 3' homology his3D1 leu2D0 lys2D0 gal80::KANMX gal4::spHIS5 | pRPL18B-eMagA, pTEF1-eMagBM | This work |

| | | | | |
|---|---|---|---|---|
| yMM1771 | pHHF1-Gal4DBD-eMagA, pHHF1-eMagBM-Gal4AD, pGAL1-mScarlet-I | BY Matα ura3Δ0::5' Ura3 homology, pHHF1-Gal4DBD-eMagA-tENO1, pHHF1-eMagBM-Gal4AD-tENO1, pGAL1-mScarlet-I-tENO1, Ura3, Ura 3' homology his3D1 leu2D0 lys2D0 gal80::KANMX gal4::spHIS5 | pHHF1-eMagA, pHHF1-eMagBM | This work |
| yMM1772 | pHHF1-Gal4DBD-eMagA, pTEF1-eMagBM-Gal4AD, pGAL1-mScarlet-I | BY Matα ura3Δ0::5' Ura3 homology, pHHF1-Gal4DBD-eMagA-tENO1, pTEF1-eMagBM-Gal4AD-tENO1, pGAL1-mScarlet-I-tENO1, Ura3, Ura 3' homology his3D1 leu2D0 lys2D0 gal80::KANMX gal4::spHIS5 | pHHF1-eMagA, pTEF1-eMagBM | This work |
| yMM1773 | pTEF1-Gal4DBD-eMagA, pTEF1-eMagBM-Gal4AD, pGAL1-mScarlet-I | BY Matα ura3Δ0::5' Ura3 homology, pTEF1-Gal4DBD-eMagA-tENO1, pTEF1-eMagBM-Gal4AD-tENO1, pGAL1-mScarlet-I-tENO1, Ura3, Ura 3' homology his3D1 leu2D0 lys2D0 gal80::KANMX gal4::spHIS6 | pTEF1-eMagA, pTEF1-eMagBM | This work |
| yMM1774 | pRPL18B-Gal4DBD-eMagA, pHHF1-eMagB-Gal4AD, pGAL1-mScarlet-I | BY Matα ura3Δ0::5' Ura3 homology, pRPL18B-Gal4DBD-eMagA-tENO1, pHHF1-eMagB-Gal4AD-tENO1, pGAL1-mScarlet-I-tENO1, Ura3, Ura 3' homology his3D1 leu2D0 lys2D0 gal80::KANMX gal4::spHIS6 | pRPL18B-eMagA, pHHF1-eMagB | This work |
| yMM1775 | pHHF1-Gal4DBD-eMagA, pHHF1-eMagB-Gal4AD, pGAL1-mScarlet-I | BY Matα ura3Δ0::5' Ura3 homology, pHHF1-Gal4DBD-eMagA-tENO1, pHHF1-eMagB-Gal4AD-tENO1, pGAL1-mScarlet-I-tENO1, Ura3, Ura 3' homology his3D1 leu2D0 lys2D0 gal80::KANMX gal4::spHIS6 | pHHF1-eMagA, pHHF1-eMagB | This work |
| yMM1776 | pHHF1-Gal4DBD-eMagA, pTEF1-eMagB-Gal4AD, pGAL1-mScarlet-I | BY Matα ura3Δ0::5' Ura3 homology, pHHF1-Gal4DBD-eMagA-tENO1, pTEF1-eMagB-Gal4AD-tENO1, pGAL1-mScarlet-I-tENO1, Ura3, Ura 3' homology his3D1 leu2D0 lys2D0 gal80::KANMX gal4::spHIS6 | pHHF1-eMagA, pTEF1-eMagB | This work |
| yMM1778 | pRPL18B-Gal4DBD-eMagAF, pRPL18B-eMagB-Gal4AD, pGAL1-mScarlet-I | BY Matα ura3Δ0::5' Ura3 homology, pRPL18B-Gal4DBD-eMagAF-tENO1, pRPL18B-eMagB-Gal4AD-tENO1, pGAL1-mScarlet-I-tENO1, Ura3, Ura 3' homology his3D1 leu2D0 lys2D0 gal80::KANMX gal4::spHIS7 | pRPL18B-eMagAF, pRPL18B-eMagB | This work |

#### Supplemental Table 2: Primers used in this study

| ID | Alias | Sequence | Target | Purpose |
|---|---|---|---|---|
| oMM2093 | Fwd NLS-GAL4AD-CIB1/N/81 3 | GCATCGTCTCATCGGTCTCATATGgataaagcggaattaattccc | NLS-GAL4AD-CIB1 | Amplify NLS-GAL4AD-CIB1 from Addgene #28245 (pMM0159) to insert into a part 3 vector |
| oMM2094 | Rev NLS-GAL4AD-CIB1/N 3 | ATGCCGTCTCAGGTCTCAGGATccgtatctacgattcatctgcagc | NLS-GAL4AD-CIB1 | Amplify NLS-GAL4AD-CIB1 from Addgene #28245 (pMM0159) to insert into a part 3 vector. Adds a GG |
| oMM2096 | Fwd GAL4BD 3a | GCATCGTCTCATCGGTCTCATATGaagctactgtcttctatcg | GAL4DBD | Amplify GAL4DBD from Addgene #28244 (pMM0212) to insert into a part 3a vector |
| oMM2097 | Rev GAL4BD 3a | ATGCCGTCTCAGGTCTCAAGAAcccgatacagtcaactgtctttg | GAL4DBD | Amplify GAL4DBD from Addgene #28244 (pMM0212) to insert into a part 3a vector. Adds a GG. |
| oMM2098 | Fwd -CRY2/PHR/515/535 3b | GCATCGTCTCATCGGTCTCATTCTacaggtgctagcttcatg | -CRY2/PHR/535 | Amplify -CRY2, PHR, 535 from pMM0521 to insert into a part 3b vector |
| oMM2100 | Rev -CRY2(535) 3b | ATGCCGTCTCAGGTCTCAGGATCCaacagccgaaggtacttg | -CRY2(535) | Amplify -CRY2(535) from pMM0521 to insert into a part 3b vector. Adds |

| | | | | |
|---|---|---|---|---|
| | | | | a GG |
| oMM2130 | Fwd VVD-Gal4AD 3 part | GCATCGTCTCATCGGTCTCATatgAGCCATACCGTGAACTC | VVD-Gal4AD | Amplify VVD-Gal4AD from pRS425-VVD to make a type 3B part |
| oMM2131 | Rev VVD-Gal4AD 3 part | ATGCCGTCTCAGGTCTCAGGATccTGAACCAGCGTAATCTGGTAC | VVD-Gal4AD | Amplify VVD-Gal4AD from pRS425-VVD to make a type 3B part. Adds a GG |
| oMM2675 | Fwd mScarlet-I 3 part | gcatCGTCTCaTCGGTCTCAtatgATGGTAAGCAAAGGTGAGG | mScarlet-I | Amplify mScarlet-I from pMM0610 to make a type 3 part |
| oMM2676 | Rev mScarlet-I 3 part | atgcCGTCTCaGGTCTCaggatCTACTTATATAATTCGTCCATGCC | mScarlet-I | Amplify mScarlet-I from pMM0610 to make a type 3 part |
| oMM2677 | Fwd Gal4AD c-term 3B | gcatCGTCTCaTCGGTCTCAttctGATAAAGCGGAATTAATTCCCG | Gal4AD | Amplify Gal4AD from pMM0928 to make a type 3B part |
| oMM2678 | Rev Gal4AD c-term 3B | atgcCGTCTCaGGTCTCaggatTTACTCTTTTTTTGGGTTTGGTGG | Gal4AD | Amplify Gal4AD from pMM0928 to make a type 3B part |
| oMM2683 | Fwd eMagB Q5 Y99F | AACACTATCTtCACCATGAAGAAGG | eMagB | Q5 to make eMagBF (Y99F) |
| oMM2684 | Rev eMagB Q5 Y99F | GGAGTCCACGTATTTGCG | eMagB | Q5 to make eMagBF (Y99F) |
| oMM2685 | Fwd eMagA Q5 Y99F | AACACGATCTtCACCATCAAG | eMagA | Q5 to make eMagAF (Y99F) |

| | | | | |
|---|---|---|---|---|
| oMM2686 | Rev eMagA Q5 Y99F | CGAGTCCACA TATTTGCG | eMagA | Q5 to make eMagAF (Y99F) |
| oMM2687 | Fwd eMagB backbone | GGCGAATACC GGTACAGCAT C | eMagB | Make eMagB backbone for Gibson |
| oMM2688 | Rev eMagB backbone(I) | aggggtgtccttctgc ttaaggtcacacag GATgagggcgcag gagaggtc | eMagB | Make eMagB backbone (I-) for Gibson |
| oMM2689 | Fwd eMagB -VMM insert | ttaagcagaaggac acccctgtg | eMagB | Make eMagB -VMM insert for Gibson |
| oMM2691 | Rev eMagB insert | GATGCTGTAC CGGTATTCGC C | eMagB | Make eMagB insert for Gibson |

#### Supplemental Table 3: Plasmids used in this study

| ID | Alias | Gene(s) or Insert | Yeast Marker | Bacterial Resistance | Toolkit Part | Source |
|---|---|---|---|---|---|---|
| pMM0159 | pGal4AD-CIB1 (Addgene #28245) | GAL4AD-CIB1 | LEU2 | AmpR | N/A | Kennedy et al. 2010 Nature Methods |
| pMM0212 | pGal4DBD-CRY2PHR (Addgene #28244) | GAL4BD-CRY2PHR TRP1 | TRP1 | KanR | N/A | Kennedy et al. 2010 Nature Methods |
| pMM0452 | Entry vector (pYTK001; Addgene #65108) | ColE1-CamR-sfGFP dropout | | CamR | Entry vector | Lee et al. 2015 Acs Syn Bio |
| pMM0454 | pRPL18B (pYTK017; Addgene #65124) | ColE1-CamR-pRPL18B | | CamR | Part, Type 2 | Lee et al. 2015 Acs Syn Bio |

| | | | | | | |
|---|---|---|---|---|---|---|
| pMM0458 | GFP dropout cassette (pYTK096) | ColE1-KanR-URA3 3' homology-URA3-sfGFP dropout-URA3 5' homology | URA3 | KanR | GFP dropout cassette | Lee et al. 2015 Acs Syn Bio |
| pMM0479 | LEU2 (pYTK075; Addgene #65182) | ColE1-CamR-LEU2 | | CamR | Part, Type 6 | Lee et al. 2015 Acs Syn Bio |
| pMM0489 | ConLS (pYTK002; Addgene #65109) | ColE1-CamR-ConLS | | CamR | Part, Type 1 | Lee et al. 2015 Acs Syn Bio |
| pMM0491 | ConR1 (pYTK067; Addgene #65174) | ColE1-CamR-ConR1 | | CamR | Part, Type 5 | Lee et al. 2015 Acs Syn Bio |
| pMM0521 | NLS-ZIF268 DBD-CRY2 (L3) | SV40NLS-Zif268DBD-CRY2 (L3) scTRP1 | TRP1 | AmpR | N/A | An-adirekkun et al. 2020 Biotechnol Bioeng |
| pMM0522 | pTEF1 (pYTK013; Addgene #65120) | ColE1-CamR-pTEF1 | | CamR | Part, Type 2 | Lee et al. 2015 Acs Syn Bio |
| pMM0524 | CEN6/ARS4 (pYTK081; Addgene #65188) | ColE1-CamR-CEN6/ARS4 | | CamR | Part, Type 7 | Lee et al. 2015 Acs Syn Bio |
| pMM0525 | RFP (pYTK084; Addgene #65191) | ColE1-KanR-RFP | | KanR | Part, Type 8 | Lee et al. 2015 Acs Syn Bio |
| pMM0532 | ConL1 (pYTK003; Addgene #65110) | ColE1-CamR-ConL1 | | CamR | Part, Type 1 | Lee et al. 2015 Acs Syn Bio |
| pMM0533 | ConL2 (pYTK004; Addgene #65111) | ColE1-CamR-ConL2 | | CamR | Part, Type 1 | Lee et al. 2015 Acs Syn Bio |

| | | | | | | |
|---|---|---|---|---|---|---|
| pMM0537 | ConR2 (pYTK068; Addgene #65175) | ColE1-CamR-ConR2 | | CamR | Part, Type 5 | Lee et al. 2015 Acs Syn Bio |
| pMM0541 | ConRE (pYTK072; Addgene #65179) | ColE1-CamR-ConRE | | CamR | Part, Type 5 | Lee et al. 2015 Acs Syn Bio |
| pMM0542 | tENO1 (pYTK051; Addgene #65158) | ColE1-CamR-tENO1 | | CamR | Part, Type 4 | Lee et al. 2015 Acs Syn Bio |
| pMM0547 | Spacer (pYTK048; Addgene #65155) | ColE1-CamR-Spacer | | CamR | Spacer | Lee et al. 2015 Acs Syn Bio |
| pMM0556 | sfGFP dropout (pYTK095; Addgene #65202) | ColE1-AmpR-sfGFP dropout | | AmpR | 678 | Lee et al. 2015 Acs Syn Bio |
| pMM0610 | mScarlet-I | mSCarlet-I caURA3 | URA3 | AmpR | N/A | Bindels et al. 2016 Nature Methods |
| pMM0773 | Spacer | ConLS-Spacer-ConR1-Amp/Cole1 | | AmpR | Cassette | This work |
| pMM0777 | Spacer | ConL1-Spacer-ConR2-Leu2-CEN/ARS-ColE1-KanR | | AmpR | Cassette | Geller et al. 2019 Cell Mol Bioeng |
| pMM0916 | GAL4AD-CIB1 | GAL4AD-CIB1 (domesticated) | LEU2 | AmpR | | This work |
| pMM0918 | Gal4DBD (n-terminal) | ColE1-CamR-Gal4DBD | | CamR | Part, Type 3a | This work |
| pMM0920 | CRY2(535) | ColE1-CamR-CRY2(535) | | CamR | Part, Type 3b | This work |
| pMM0921 | CRY2PHR | ColE1-CamR-CRY2PHR | | CamR | Part, Type 3b | This work |

| | | | | | | |
|---|---|---|---|---|---|---|
| pMM0922 | CRY2FL | ColE1-CamR-CRY2FL | | CamR | Part, Type 3b | This work |
| pMM0923 | Gal4AD-CIB1 | ColE1-CamR-Gal4AD-CIB1 | | CamR | Part, Type 3 | This work |
| pMM0928 | VVD-Gal4AD | ColE1-CamR-VVD-Gal4AD | | CamR | Part, Type 3 | This work |
| pMM0939 | pGAL1 (pYTK030; #Addgene #65137) | ColE1-CamR-pGAL1 | | CamR | Part, Type 2 | Lee et al. 2015 Acs Syn Bio |
| pMM0943 | pRPL18B-Gal4DBD-CRY2(535) cassette | ConLS-pRPL18B-Gal4DBD-CRY2(535)-tENO1-ConR1 | | AmpR | Cassette | This work |
| pMM0944 | pRPL18B-Gal4DBD-CRY2PHR cassette | ConLS-pRPL18B-Gal4DBD-CRY2PHR-tENO1-ConR1 | | AmpR | Cassette | This work |
| pMM0945 | pRPL18B-Gal4DBD-CRY2FL cassette | ConLS-pRPL18B-Gal4DBD-CRY2FL-tENO1-ConR1 | | AmpR | Cassette | This work |
| pMM0946 | pRPL18B-Gal4AD-CIB1 cassette | ConL1-pRPL18B-Gal4AD-CIB1-tENO1-ConR2 | | AmpR | Cassette | This work |
| pMM1104 | pHHF1 (pYTK012; #Addgene #65119) | ColE1-CamR-pHHF1 | | CamR | Part, Type 2 | Lee et al. 2015 Acs Syn Bio |
| pMM1235 | mScarlet-I | ColE1-CamR-mScarlet-I | | CamR | Part, Type 3 | This work |
| pMM1236 | pGAL1-mScarlet cassette | ConL2-pGAL1-mScarlet-I-tENO1-ConRE | | AmpR | Cassette | This work |
| pMM1237 | pRPL18B-mScarlet-I cassette | ConL2-pRPL18B-mScarlet-I-tENO1-ConRE | | AmpR | Cassette | This work |
| pMM1239 | spacer, spacer, | 5' Ura3 homology, spacer, spacer, | URA3 | KanR | Multigene | This work |

| | | | | | | |
|---|---|---|---|---|---|---|
| | pGAL1-mScarlet-I | pGAL1-mScarlet-I-tENO1, Ura3, Ura 3' homology KanR-ColE1 | | | | |
| pMM1240 | spacer, spacer, pRPL18B-mScarlet-I multigene | 5' Ura3 homology, spacer, spacer, pRPL18B-mScarlet-I-tENO1, Ura3, Ura 3' homology KanR-ColE1 | URA3 | KanR | Multigene | This work |
| pMM1242 | pRPL18B-Gal4DBD-CRY2PHR, pRPL18B-Gal4AD-CIB1, pGAL1-mScarlet-I | 5' Ura3 homology, pRPL18B-Gal4DBD-CRY2PHR-tENO1, pRPL18B-Gal4AD-CIB1-tENO1, pGAL1-mScarlet-I-tENO1, Ura3, Ura 3' homology KanR-ColE1 | URA3 | KanR | Multigene | This work |
| pMM1244 | pRPL18B-Gal4DBD-CRY2FL, pRPL18B-Gal4AD-CIB1, pGAL1-mScarlet-I | 5' Ura3 homology, pRPL18B-Gal4DBD-CRY2FL-tENO1, pRPL18B-Gal4AD-CIB1-tENO1, pGAL1-mScarlet-I-tENO1, Ura3, Ura 3' homology KanR-ColE1 | URA3 | KanR | Multigene | This work |
| pMM1245 | eMagB | ColE1-CamR-eMagB | | CamR | Part, Type 3a | This work |
| pMM1246 | eMagA | ColE1-CamR-eMagA | | CamR | Part, Type 3b | This work |
| pMM1247 | Gal4AD | ColE1-CamR-Gal4AD | | CamR | Part, Type 3b | This work |
| pMM1248 | pRPL18B-Gal4DBD-eMagA | ConLS-pRPL18B-Gal4DBD-eMagA-tENO1-ConR1 | | AmpR | Cassette | This work |
| pMM1249 | ConL1-pRPL18B-eMagB-Gal4AD | ConL1-pRPL18B-eMagB-Gal4AD-tENO1-ConR2 | | AmpR | Cassette | This work |
| pMM1250 | pRPL18B-Gal4DBD-eMa | 5' Ura3 homology, pRPL18B-Gal4DBD- | URA3 | KanR | Multigene | This work |

| | | | | | | |
|---|---|---|---|---|---|---|
| | gA, pRPL18B-eMagB-AD, pGAL1-mScarlet-I | eMagA-tENO1, pRPL18B-eMagB-AD-tENO1, pGAL1-mScarlet-I-tENO1, Ura3, Ura 3' homology KanR-ColE1 | | | | |
| pMM1251 | pTEF1-Gal4DBD-eMagA | ConLS-pTEF1-Gal4DBD-eMagA-tENO1-ConR1 | | AmpR | Cassette | This work |
| pMM1252 | pTEF1-eMagB-Gal4AD | ConL1-pTEF1-eMagB-Gal4AD-tENO1-ConR2 | | AmpR | Cassette | This work |
| pMM1253 | pRPL18B-Gal4DBD-eMagA, pTEF1-eMagB-AD, pGAL1-mScarlet-I | 5' Ura3 homology, pRPL18B-Gal4DBD-eMagA-tENO1, pTEF1-eMagB-AD-tENO1, pGAL1-mScarlet-I-tENO1, Ura3, Ura 3' homology KanR-ColE1 | URA3 | KanR | Multigene | This work |
| pMM1254 | pTEF1-Gal4DBD-eMagA, pTEF1-eMagB-AD, pGAL1-mScarlet-I | 5' Ura3 homology, pTEF1-Gal4DBD-eMagA-tENO1, pTEF1-eMagB-AD-tENO1, pGAL1-mScarlet-I-tENO1, Ura3, Ura 3' homology KanR-ColE1 | URA3 | KanR | Multigene | This work |
| pMM1257 | Gal4DBD-CRY2(535) Gal4AD-CIB1, pGAL1-mScarlet-I | 5' Ura3 homology, pRPL18B-Gal4DBD-CRY2(535)-tENO1, pRPL18B-Gal4AD-CIB1-tENO1, pGAL1-mScarlet-I-tENO1, Ura3, Ura 3' homology KanR-ColE1 | URA3 | KanR | Multigene | This work |
| pMM1258 | eMagBF | ColE1-CamR-eMagBF | | CamR | Part, Type 3a | This work |
| pMM1259 | eMagAF | ColE1-CamR-eMagAF | | CamR | Part, Type 3b | This work |

| | | | | | | |
|---|---|---|---|---|---|---|
| pMM1260 | eMagBM | ColE1-CamR-eMagBM | | CamR | Part, Type 3a | This work |
| pMM1263 | pRPL18B-Gal4DBD-eMagAF | ConLS-pRPL18B-Gal4DBD-eMagAF-tENO1-ConR1 | | AmpR | Cassette | This work |
| pMM1264 | pRPL18B-eMagBF-Gal4AD | ConL1-pRPL18B-eMagBF-Gal4AD-tENO1-ConR2 | | AmpR | Cassette | This work |
| pMM1265 | pRPL18B-eMagBM-Gal4AD | ConL1-pRPL18B-eMagBM-Gal4AD-tENO1-ConR2 | | AmpR | Cassette | This work |
| pMM1269 | pRPL18B-Gal4DBD-eMagAF-tENO1, pRPL18B-eMagBF-Gal4AD-tENO1, pGAL1-mScarlet | 5' Ura3 homology, pRPL18B-Gal4DBD-eMagAF-tENO1, pRPL18B-eMagBF-Gal4AD-tENO1, pGAL1-mScarlet-I-tENO1, Ura3, Ura 3' homology KanR-ColE1 | URA3 | KanR | Multigene | This work |
| pMM1270 | pRPL18B-Gal4DBD-eMagA-tENO1, pRPL18B-eMagBM-Gal4AD-tENO1, pGAL1-mScarlet-I | 5' Ura3 homology, pRPL18B-Gal4DBD-eMagA-tENO1, pRPL18B-eMagBM-Gal4AD-tENO1, pGAL1-mScarlet-I-tENO1, Ura3, Ura 3' homology KanR-ColE1 | URA3 | KanR | Multigene | This work |
| pMM1315 | pHHF1-Gal4DBD-eMagA | ConLS-pHHF1-Gal4DBD-eMagA-tENO1-ConR1 | | AmpR | Cassette | This work |
| pMM1316 | pHHF1-eMagBM-Gal4AD | ConL1-pHHF1-eMagBM-Gal4AD-tENO1-ConR2 | | AmpR | Cassette | This work |
| pMM1317 | pTEF1-eMagBM-Gal4AD | ConL1-pTEF1-eMagBM-Gal4AD-tENO1-ConR2 | | AmpR | Cassette | This work |
| pMM1318 | pHHF1-eMagB-Gal4AD | ConL1-pHHF1-eMagB-Gal4AD-tENO1-ConR2 | | AmpR | Cassette | This work |

| | | | | | | |
|---|---|---|---|---|---|---|
| pMM1319 | pRPL18B-Gal4DBD-eMagA-tENO1, pHHF1-eMagBM-Gal4AD-tENO1, pGAL1-mScarlet-I | 5' Ura3 homology, pRPL18B-Gal4DBD-eMagA-tENO1, pHHF1-eMagBM-Gal4AD-tENO1, pGAL1-mScarlet-I-tENO1, Ura3, Ura 3' homology KanR-ColE1 | URA3 | KanR | Multigene | This work |
| pMM1320 | pRPL18B-Gal4DBD-eMagA-tENO1, pTEF1-eMagBM-Gal4AD-tENO1, pGAL1-mScarlet-I | 5' Ura3 homology, pRPL18B-Gal4DBD-eMagA-tENO1, pTEF1-eMagBM-Gal4AD-tENO1, pGAL1-mScarlet-I-tENO1, Ura3, Ura 3' homology KanR-ColE1 | URA3 | KanR | Multigene | This work |
| pMM1321 | pHHF1-Gal4DBD-eMagA-tENO1, pHHF1-eMagBM-Gal4AD-tENO1, pGAL1-mScarlet- | 5' Ura3 homology, pHHF1-Gal4DBD-eMagA-tENO1, pHHF1-eMagBM-Gal4AD-tENO1, pGAL1-mScarlet-I-tENO1, Ura3, Ura 3' homology KanR-ColE1 | URA3 | KanR | Multigene | This work |
| pMM1322 | pHHF1-Gal4DBD-eMagA-tENO1, pTEF1-eMagBM-Gal4AD-tENO1, pGAL1-mScarlet-I | 5' Ura3 homology, pHHF1-Gal4DBD-eMagA-tENO1, pTEF1-eMagBM-Gal4AD-tENO1, pGAL1-mScarlet-I-tENO1, Ura3, Ura 3' homology KanR-ColE1 | URA3 | KanR | Multigene | This work |
| pMM1323 | pTEF1-Gal4DBD-eMagA-tENO1, pTEF1-eMagBM-Gal4AD-tENO1, pGAL1-mScarlet-I | 5' Ura3 homology, pTEF1-Gal4DBD-eMagA-tENO1, pTEF1-eMagBM-Gal4AD-tENO1, pGAL1-mScarlet-I-tENO1, Ura3, Ura 3' homology KanR-ColE1 | URA3 | KanR | Multigene | This work |

| | | | | | | |
|---|---|---|---|---|---|---|
| pMM1324 | pRPL18B-Gal4DBD-eMagA-tENO1, pHHF1-eMagB-Gal4AD-tENO1, pGAL1-mScarlet-I-tENO1 | 5' Ura3 homology, pRPL18B-Gal4DBD-eMagA-tENO1, pHHF1-eMagB-Gal4AD-tENO1, pGAL1-mScarlet-I-tENO1, Ura3, Ura 3' homology KanR-ColE1 | URA3 | KanR | Multigene | This work |
| pMM1325 | pHHF1-Gal4DBD-eMagA-tENO1, pHHF1-eMagB-Gal4AD-tENO1, pGAL1-mScarlet-I | 5' Ura3 homology, pHHF1-Gal4DBD-eMagA-tENO1, pHHF1-eMagB-Gal4AD-tENO1, pGAL1-mScarlet-I-tENO1, Ura3, Ura 3' homology KanR-ColE1 | URA3 | KanR | Multigene | This work |
| pMM1326 | pHHF1-Gal4DBD-eMagA-tENO1, pTEF1-eMagB-Gal4AD-tENO1, pGAL1-mScarlet-I-tENO1 | 5' Ura3 homology, pHHF1-Gal4DBD-eMagA-tENO1, pTEF1-eMagB-Gal4AD-tENO1, pGAL1-mScarlet-I-tENO1, Ura3, Ura 3' homology KanR-ColE1 | URA3 | KanR | Multigene | This work |
| pMM1328 | pRPL18B-Gal4DBD-eMagAF-tENO1, pRPL18B-eMagB-Gal4AD-tENO1, pGAL1-mScarlet | 5' Ura3 homology, pRPL18B-Gal4DBD-eMagAF-tENO1, pRPL18B-eMagB-Gal4AD-tENO1, pGAL1-mScarlet-I-tENO1, Ura3, Ura 3' homology KanR-ColE1 | URA3 | KanR | Multigene | This work |
| pRS425-VVD | VVD-GAL4AD | pADH1-VVD-GAL4AD-tADH2 | URA3 | AmpR | | Salinas et al. 2018 mBio |

Supplemental Table 4: Gene blocks used in this study

| ID | Alias | Sequence | Purpose |
|---|---|---|---|
| gMM035 | Gal4DBD | GCATCGTCTCATCGGTCTCATATGaagctactgtcttctatcgaacaagcatgcgatatttgccgacttaaaaagctcaagtgctccaaagaaaaaccgaagtgcgccaagtgtctgaagaacaactgggagtgtcgctactctcccaaaaccaaaaggtcaccgctgactagggcacatctgacagaagtggaatcaaggctagaaagactggaacagctatttctactgatttttcctcgagaagaccttgacatgattttgaaaatggattctttacaggatataaaagcattgttaacaggattatttgtacaagataatgtgaataaagatgccgtcacagatagattggcttcagtggagactgatatgcctctaacattgagacagcatagaataagtgcgacatcatcatcggaagagagtagtaacaaaggtcaaagacagttgactgtatcgggTTCTTGAGACCTGAGACGGCAT | Domesticated Gal4DBD with flanking regions for insertion into a type 3a yOTK part plasmid (pMM0918) |
| gMM055 | eMagB entry part 3A | gcatCGTCTCaTCGGTCTCAtatgggacataccctctacgcgccgggggggttatgacatcatgggttacctcagacagatcagaaaccggccgaacccacaagtggagctgggacccgtcgacctctcctgcgccctcgtgctgtgtgaccttaagcagaaggacacccctgtggtgtacgcctccgaagcattcctggagatgaccgggtacaacagacacgaagtgctgggacggaactgccgcttcctgcaatccccggatggaatggtgaagcctaagtcaacccgcaaatacgtggactccaacactatctacaccatgaagaaggccattgaccgcaatgctgaggtgcaagtggaagtggtgaacttcaagaagaacggacagcgcttcgtcaacttcctgactatgattcccgtgcgggacgaaaccggcgaataccggtacagcatcgggtttcagtgcgagactgagGGCGGTGGCGGCAGTGGTGGAATGAAGCAACTCGAGGACAAGGTTGAGGAACTGCTGAGTAAGAATTACCACCTCGAAAACGAGGTCGCACGATTGAAAAAGTTGGTGGGTGAGggaggtggtggatcgggtggaggTTCTTGAGACCtGAGACGgcat | Makes eMagB entry part 3A with BsmBI digestion |
| gMM056 | eMagA entry part 3B | gcatCGTCTCaTCGGTCTCAttctggaggcggaggctccggtggtggtggaagtggcggtggcggatccatgggacacactctttacgcccctggaggatacgacattatgggatatttggatcagattgcgaaccgcccaaaccctcaggtcgaactggggcctgtggacctgtcatgtgccctgatcctgtgcgatctgaagcaaaaggacactccgatcgtctacgcctcggaagccttcttggagatgaccggatacaacagacatgaggtgctcggcaggaactgcagattcctgcagtcccccgacgggatggtgaaaccaaagtcg | Makes eMagA entry part 3B with BsmBI digestion |

| | | |
|---|---|---|
| | | actcgcaaatatgtggactcgaacacgatctaca ccatcaagaaggccatcgaccggaacgccgag gtccaggtggaggtggtcaactttaagaagaacg gccagcggttcgtgaactttctgaccatcattccgg tccgggatgaaaccggagagtacagatactcca tcggattccagtgcgaaaccgaaTAAatcctGA GACCtGAGACGgcat |
